# The transcription factor RIP140 regulates interferon γ signaling in breast cancer

**DOI:** 10.1101/2024.02.15.580503

**Authors:** S. Jalaguier, A. Kuehn, C. Petitpas, A. Dulom, R. Jacquemont, C. Assi, S. Sixou, U. Jeschke, J. Colinge, V. Cavaillès

**Affiliations:** IRCM, Institut de Recherche en Cancérologie de Montpellier, Montpellier, F-34298, France; INSERM, U1194, Montpellier, F-34298, France; Université de Montpellier, Montpellier, F-34090, France; Institut régional du Cancer de Montpellier, Montpellier, F-34298, France; Faculté des Sciences Pharmaceutiques, Université Toulouse III—Paul Sabatier, F-31062 Toulouse, France; Department of Obstetrics and Gynecology, University Hospital Augsburg, Stenglinstrasse 2, 86156 Augsburg, Germany; CNRS, Montpellier, F-34298, France

**Keywords:** RIP140/NRIP1, interferon-γ, GBP1, gene expression, cell proliferation, breast cancer

## Abstract

RIP140 (receptor interacting protein of 140 kDa) is an important player in breast cancer (BC) by regulating key cellular pathways such as nuclear hormone receptors signaling. In order to identify additional genes specifically regulated by RIP140 in BC, we performed an RNA sequencing after silencing its expression in MCF-7 cells. We identified the interferon γ (IFNγ) signaling as being substantially repressed by RIP140 knock-down. Using the *GBP1* (guanylate binding protein 1) gene as a reporter of IFNγ signaling, we demonstrated its robust induction by RIP140 through an ISRE motif, leading to a significant reduction of its induction upon IFNγ treatment. Furthermore, we showed that low levels of RIP140 amplified the IFNγ-dependent inhibition of BC cell proliferation. In line with these data, reanalysis of transcriptomic data obtained in human BC samples, revealed that IFNγ levels were associated with good prognosis only for BC patients exhibiting tumors expressing low levels of RIP140, thus confirming its effect on the anti-tumor activity of IFNγ provided by our experimental data. Altogether, this study identifies RIP140 as a new regulator of IFNγ signaling in breast tumorigenesis.

## INTRODUCTION

Breast cancer (BC) is the most common tumor for women in the world. In 2020, according to the World Health Organization, over two million women were diagnosed with a breast cancer and more than 600 000 died of that pathology. A diversity of signaling pathways, which strongly cross-talk, cause survival and proliferation of BC cells. From a clinical point of view, the main molecular target is estrogen receptor α (ERα) which drives around 70 % of BC ^1^. These tumors also express the progesterone receptor (PR). Another important molecular target is epidermal growth factor 2 (ERBB2, formerly HER2 or HER2/neu), a transmembrane receptor amplified or overexpressed in approximately 20% of BC ^2^. Finally, a third BC subtype, triple-negative breast cancers, characterized by the lack of expression of molecular targets ERα, PR, or ERBB2 account for approximately 15% of all breast tumors ^3^.

Other pathways are implicated including the interferon-γ/STAT1 (IFNγ/STAT1) signaling which can induce tumor cell apoptosis and results in repression of BC cell growth ^4^. In addition to prompt apoptosis and senescence, IFNγ can shift tumors to a dormant state ^5^. It must also be underlined that this pathway is believed to be critical for the success of immune therapy ^6^. However, IFNγ displays a yin and yang effect, involving repression of tumor growth by arresting cell cycle, induction of tumor ischemia, inhibition of suppressive immune cells as well as promotion of carcinogenesis and angiogenesis ^7^.

RIP140 (receptor interacting protein of 140kDa) ^8^, also known as NRIP1 (nuclear receptor interacting protein 1), was shown to play an active role in many physiological or pathological processes. For instance, mice lacking the *Rip140* gene displayed a female infertility, a reduced body fat content and severe cognitive defects ^9^. RIP140 regulates mammary gland development by directly regulating the expression of the progesterone receptor, amphiregulin and signal transducer and activator of transcription 5a ^10^. In BC, we showed that a decreased expression of RIP140 activity was associated with the bad prognosis basal-like tumors ^11^ while an increase in RIP140 mRNA was observed in overall breast tumors ^12^. Very recently, our team evidenced that RIP140 could repress proliferation of BC cells by targeting glycolysis (more precisely GLUT3 expression) ^13^ and the phosphate pentose pathway through inhibition of glucose 6-phosphate dehydrogenase ^14^.

In the present study, for a better understanding of RIP140 action in BC, we performed an RNA sequencing analysis of MCF7 cells presenting a downregulation of *NRIP1*. We herein evidenced a large number of target genes upregulated by RIP140, as expected from its role as a transcriptional repressor, but others were significantly down-regulated including for instance the vitamin D receptor. We documented the interferon signaling as a key target of RIP140 and further focused on GBP1, a main IFNγ-induced gene. We demonstrated that its IFNγ-dependent expression in three BC cell lines (MCF7, T47D and MDA-MB-231) was inhibited by RIP140. We showed that the regulation occured at the transcriptional level through an IFN-response element and that reducing RIP140 levels in BC cells amplified the IFNγ response on gene expression and cell proliferation. Finally, reanalysis of transcriptomics data from BC patient samples validated the impact of low RIP140 levels on the prognosis value of the *IFNG* gene expression. Altogether, our data demonstrated that RIP140 acts as a key regulator of the IFNγ response in BC cells.

## MATERIALS AND METHODS

### Materials

Estradiol and Interferon γ were purchased from Sigma-Aldrich ®.

### Plasmids

pEFcmyc-RIP140 was previously described ^15^. Pro3757-hGBP1, pro237-hGBP1 and ΔISRE-hGBP1 were kindly provided by Dr Elisabeth Naschberger ^16^. pISRE-F-luc was a generous gift from Dr Konstantin Maria Johannes Sparrer ^17^.

### RNA-seq data generation and processing

RNA extracts were submitted to Illumina sequencing following a single end 1×75bp protocol. Read sequences were cut to eliminate the first 13 nucleotides due to compositional bias. Low quality ends were further trimmed with sickle. The resulting shortened reads were aligned against the human genome (hg38) with the TopHat 2.1.0 algorithm. Gene expression data (read counts) were obtained applying HTSeq-count to extract the number of reads aligned over each protein-coding gene. Read counts were compiled in a gene expression matrix. Genes that did not reach a minimum read count of 5 in at least 4 samples were considered not expressed and thus discarded. The remaining expression matrix was normalized with edgeR ^18^ TMM method. We then constructed a statistical model of the whole data set using edgeR robust negative binomial generalized linear model with a design matrix linking the two replicates of each condition. Pairwise gene selections, e.g. siRIP140 against siCtl unstimulated (ethanol), were subsequently obtained from the globally calibrated model. In every case, we imposed a p-value<0.01, a minimum log_2_-fold-change of 0.5 in absolute value (1.4 in the linear scale), and an average minimum read count over all the samples of the two tested conditions of 20.

### Pathway analysis

Gene set enrichment analysis was conducted with R BioConductor FGSEA library ^19^ using GO Biological Process terms and Reactome pathways as gene sets. FGSEA was run with 1,000,000 permutations and we imposed minimum and maximum gene set sizes of 15 and 250 respectively.

### Cell culture and transfections

MELN (RRID:CVCL_WI19) cells were cultured in phenol red-free DMEM-F12 (Invitrogen) containing 5% dextran-charcoal treated FCS (Invitrogen), MCF7 (RRID:CVCL_0031) and MDA-MB-231 (RRID:CVCL_0062) in DMEM-F12 (Invitrogen) and T47D in RPMI (Invitrogen) supplemented with 10% fetal calf serum (FCS) and antibiotics (Gibco). All cell lines were authenticated by short tandem repeat profiling and free from mycoplasma. When needed, plasmids or siRNAs were transfected using either JetPEI or INTERFERin (Polyplus), respectively. When indicated, cells were treated with 100 nM E2 or 10 µg/ml IFNγ and then harvested 24 hours later. The list and sequences of the siRNAs used in this study are indicated in Supplementary table 1. SiCtl is control siRNA, designed not to target any gene.

### mRNA quantification

RNA was isolated using the Zymo Research kit (Zymo research) and reverse transcription (RT)-qPCR assays were done using qScript (VWR) according to the manufacturer’s protocol. Transcripts were quantified using SensiFAST SYBR (BioLine) on a LC480 instrument. The nucleotide sequences of the primers used for real time PCR are indicated in Supplementary Table 1.

### Immunoblotting assays

Western blot analyses were performed as previously described ^12^ using anti-VDR (Ozyme), anti-GBP1 (Origene) or anti-actin (Sigma) antibodies.

### Luciferase assays

MCF7, MDA-MB-231 and T47D cells were plated in 96-well plates 24 hours prior to transfection with either JetPEI or INTERFERin according to the manufacturer’s protocol. Firefly luciferase data were normalized to renilla luciferase activity and expressed as relative luciferase activities.

### Cell proliferation assay

MCF7, MDA-MB-231 and T47D cells were seeded at a density of 2000 cells per well. At the indicated time, 0.5 mg/ml 3-(4,5-dimethylthiazol-2-yl)-2,5-diphenyltetrazolium bromide (MTT) (Sigma-Aldrich, St Louis, MO, USA) was added to selected wells and cells were incubated at 37°C for 4h. Formazan crystals were solubilized in DMSO and absorbance read at 560 nm on a spectrophotometer. Results were normalized to the cell density at day 1.

### Survival analysis

Using the KM-plot Private Edition (http://kmplot.com) ^20^, RNA sequencing data from the Cancer Genome Atlas (TCGA) were analyzed using Cox proportional hazards regression, as previously described ^21^. The Kaplan-Meier method was used to estimate overall survival calculated from diagnosis to death. Patients lost to follow-up were censored at the time of last contact.

### Statistical analysis

All experiments were conducted independently at least three times. Results were expressed as the mean ± standard deviation (S.D.). Comparisons of two independent groups were performed using the Mann-Whitney U test. A probability level (*p* value) of 0.05 was chosen for statistical significance. Statistical analyses were performed using StatEL (AdScience).

## RESULTS

### Identification and validation of RIP140 target genes in BC cells

In order to get a comprehensive survey of RIP140 target genes in BC cells, we performed an RNA sequencing using MCF7 BC cells stably expressing luciferase under the control of estrogen responsive elements (MELN cells) ^22^. We first validated RIP140 silencing using small interfering RNA (siRIP140) transfection, at the mRNA level (Figure 1A) and at the protein level (Supplementary Figure 1A). RIP140 knock-down was also validated by a 2-fold increase of estradiol-dependent regulation of luciferase expression (Figure 1B).

**Figure 1.**
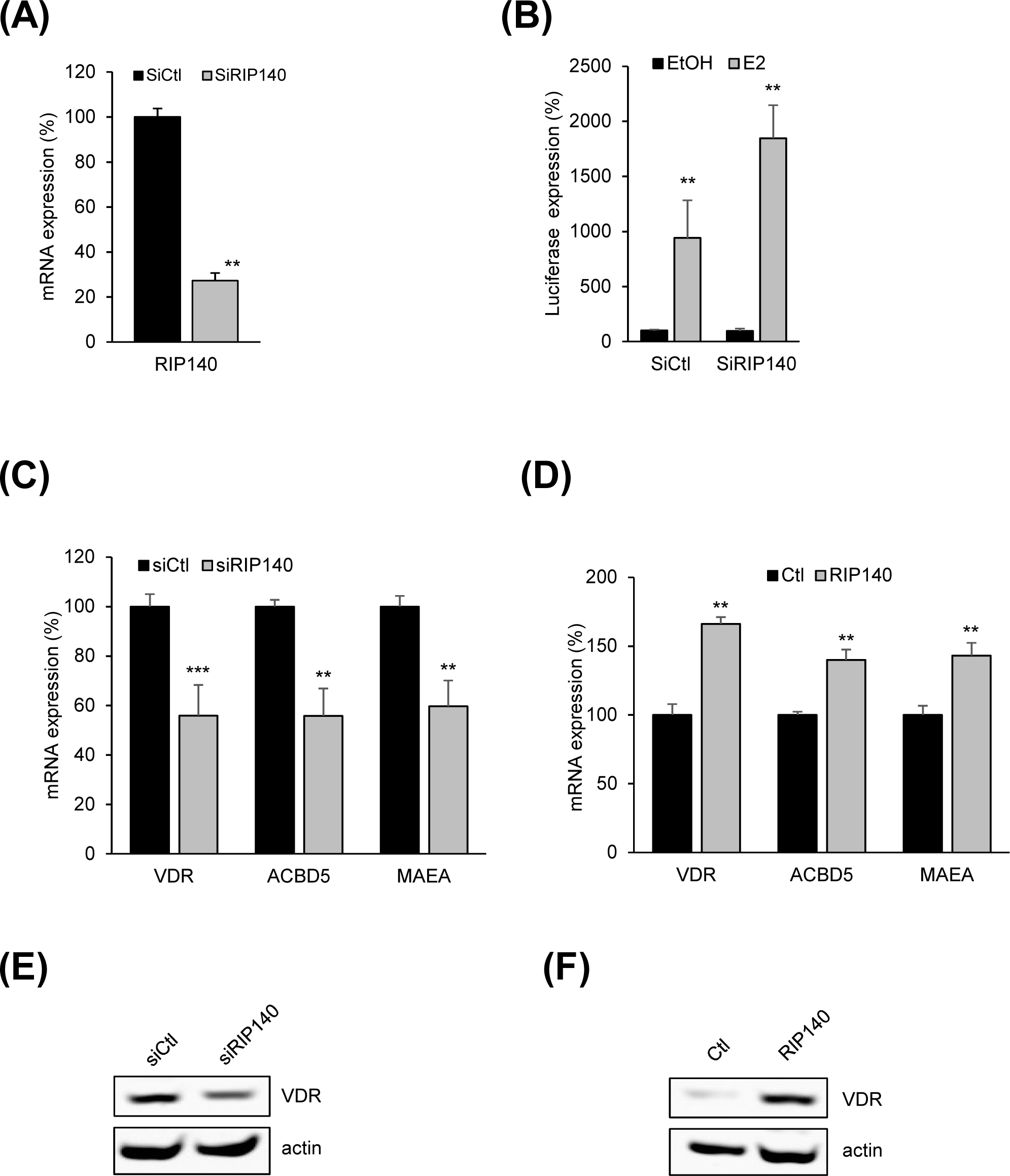
Validation of RNA sequencing results. (A) MELN cells were transfected with either control siRNA (siCtl), or siRIP140. Total RNAs were subjected to RT-qPCR assays, and quantification of RIP140 mRNA was expressed relatively to the control condition (siCtl). (B) MELN cells were transfected as in A and then treated or not with estradiol (10^-8^ M). Luciferase mRNA was quantified and results expressed as above described. MCF7 cells were transfected with siCtl or siRIP140 (C) or with pEF-RIP140 or the empty vector (D). Quantification of mRNAs was done as described in (A). (E) MCF7 cells were transfected as in (C) and western-blot assays were done with either an anti-VDR (upper panel) or an anti-actin (lower panel antibody. (F) MCF7 cells were transfected as in (D) and western-blot assays were performed as in (E). Statistical analyses were performed using the Mann-Whitney test. *p<0.05; **p<0.01; ***p<0.001.

An RNA sequencing analysis of these BC cells transfected or not with siRIP140 was then performed in the absence of estradiol. On Table 1 are shown the genes ranked according to the best p value and a negative (top panel) or positive (bottom panel) amplitude fold. Interestingly, downregulation of RIP140 expression only induced small variations of target genes level, a phenomenon we long experienced.

**Table 1.**
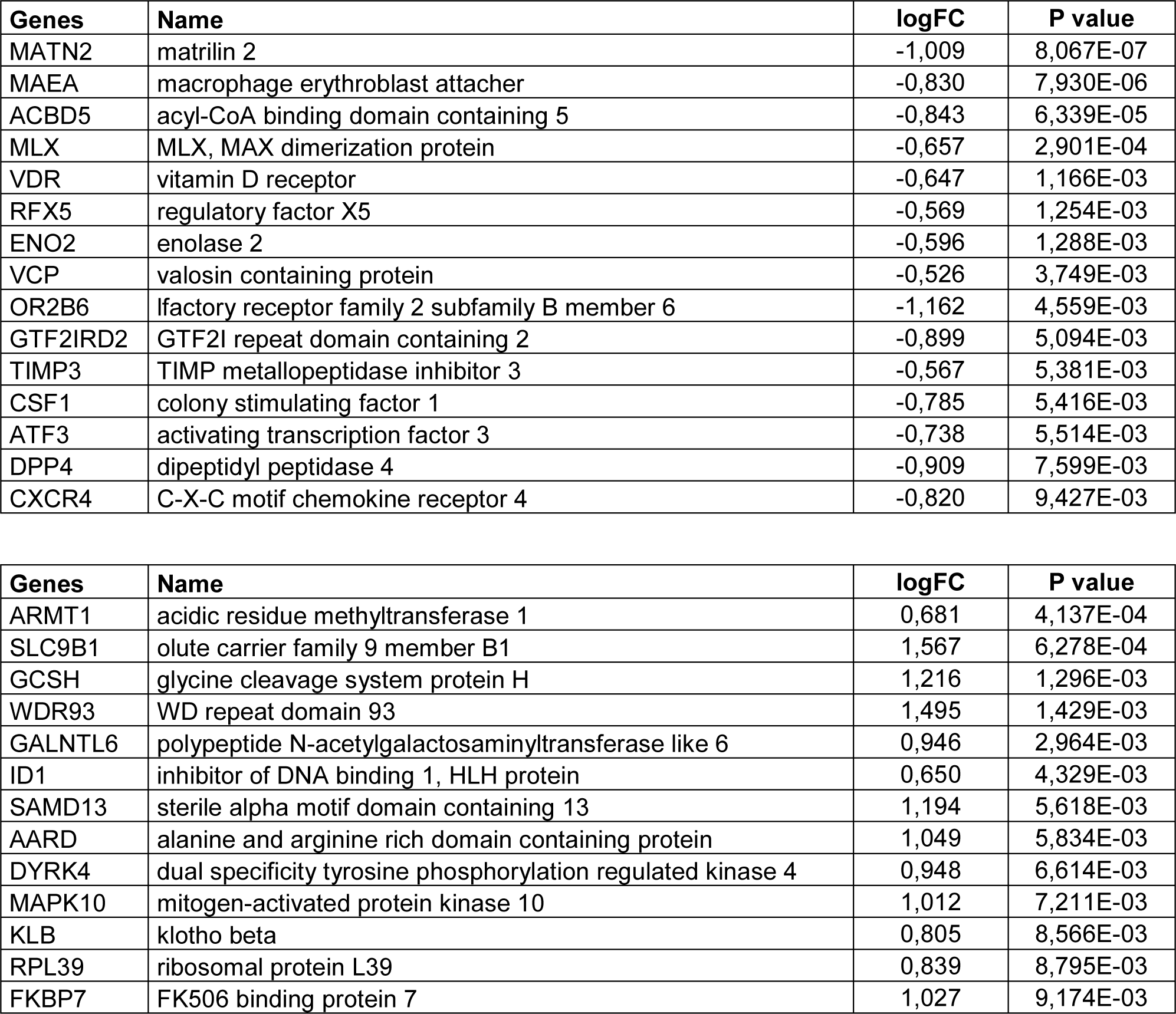
Top genes down-regulated (upper panel) or up-regulated (lower panel) after RIP140 mRNA knock-down. Genes are listed from lower to higher p values.

The RNA sequencing results were then validated by RT-qPCR assays. As shown in Figure 1C, we validated several genes down-regulated under siRIP140 transfection, including the vitamin D receptor (*VDR*), acyl-CoA binding domain containing 5 (*ACBD5*) and macrophage erythroblast attacher (*MAEA*). Ectopic expression of RIP140 in MCF7 cells, induced the opposite effect, *i.e.* an up-regulation of the three above mentioned genes (Figure 1D). Similar results were obtained in the T47D luminal BC cell line (data not shown). Concerning VDR, we also evidenced at the protein level the effect of either siRIP140 (Figure 1E) or the ectopic expression of RIP140 (Figure 1F). Altogether, these data validated the robustness of our RNA sequencing analysis.

### Interferon γ signaling is a main target of RIP140

The gene set enrichment analysis (GSEA) of our sequencing data allowed us to identify a series of over-represented genes, significantly enriched or depleted, associated to a phenotype. Accordingly, IFNγ signaling, a major pathway in breast carcinogenesis appeared on top of the list (Table 2). More precisely, five genes belonging to this pathway were identified as being regulated by RIP140, guanylate binding protein 1 (GBP1), SP100 nuclear antigen (SP100), major histocompatibility complex, class I, G (HLA-G), major histocompatibility complex, class II, DP alpha 1 (HLA-DPA1) and interferon beta 1 (IFNB) (data not shown). GBP1 being described as one of the most significantly induced proteins in cells exposed to IFNγ ^23^, and shown to play a major role in BC ^24^, this prompted us to further investigate the regulation of this gene by RIP140.

**Table 2.**
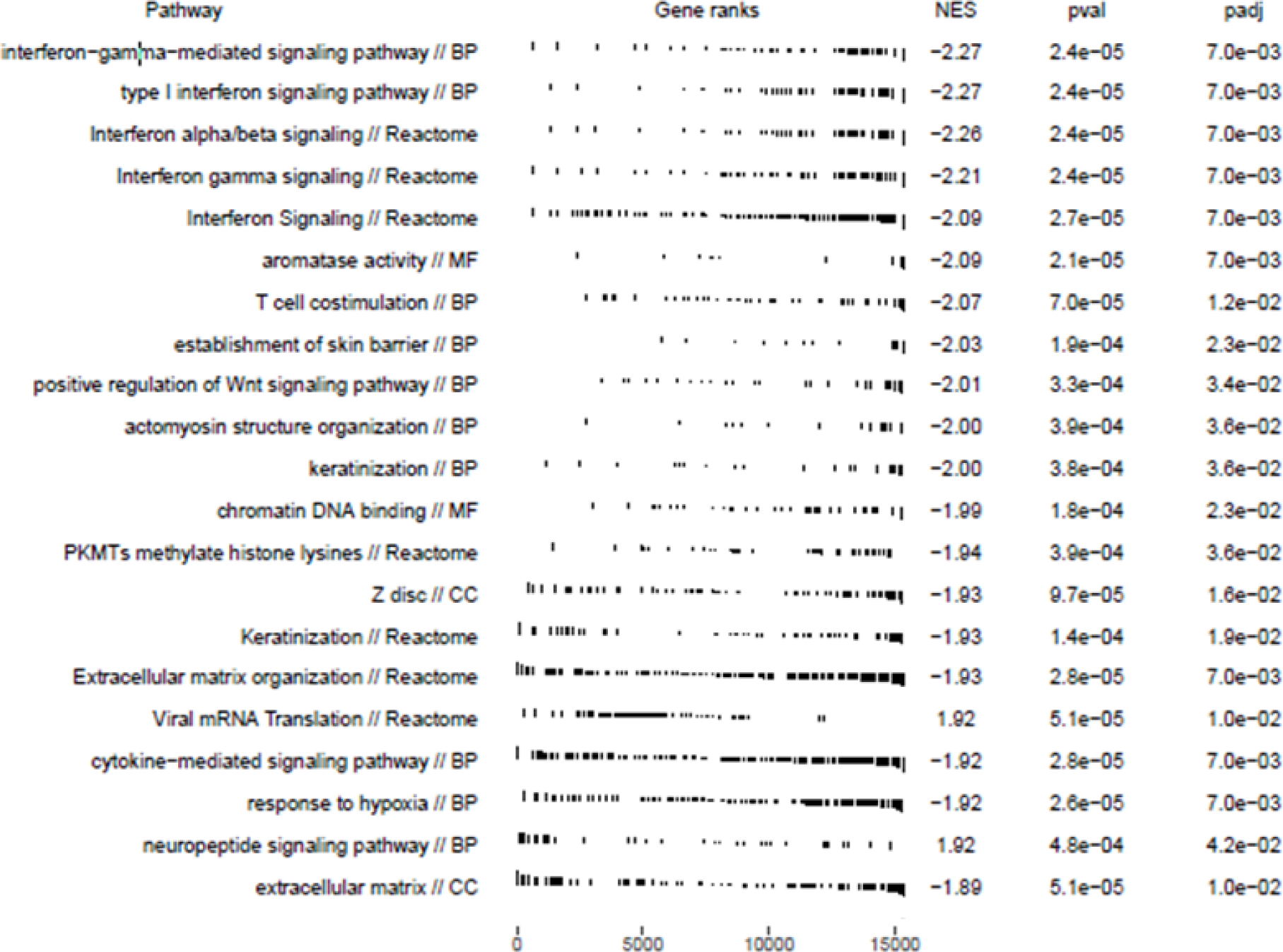
Interferon signaling is a target of RIP140. Gene set enrichment analysis of RIP140 knockdown consequences under basal conditions combining GO Biological Process terms and Reactome pathways.

| Pathway | Gene ranks | NES | pval | padj |
| --- | --- | --- | --- | --- |
| interferon-gamma-mediated signaling pathway // BP |  | -2.27 | 2.4e-05 | 7.0e-03 |
| type I interferon signaling pathway // BP |  | -2.27 | 2.4e-05 | 7.0e-03 |
| Interferon alpha/beta signaling // Reactome |  | -2.26 | 2.4e-05 | 7.0e-03 |
| Interferon gamma signaling // Reactome |  | -2.21 | 2.4e-05 | 7.0e-03 |
| Interferon Signaling // Reactome |  | -2.09 | 2.7e-05 | 7.0e-03 |
| aromatase activity // MF |  | -2.09 | 2.1e-05 | 7.0e-03 |
| T cell costimulation // BP |  | -2.07 | 7.0e-05 | 1.2e-02 |
| establishment of skin barrier // BP |  | -2.03 | 1.9e-04 | 2.3e-02 |
| positive regulation of Wnt signaling pathway // BP |  | -2.01 | 3.3e-04 | 3.4e-02 |
| actomyosin structure organization // BP |  | -2.00 | 3.9e-04 | 3.6e-02 |
| keratinization // BP |  | -2.00 | 3.8e-04 | 3.6e-02 |
| chromatin DNA binding // MF |  | -1.99 | 1.8e-04 | 2.3e-02 |
| PKMTs methylate histone lysines // Reactome |  | -1.94 | 3.9e-04 | 3.6e-02 |
| Z disc // CC |  | -1.93 | 9.7e-05 | 1.6e-02 |
| Keratinization // Reactome |  | -1.93 | 1.4e-04 | 1.9e-02 |
| Extracellular matrix organization // Reactome |  | -1.93 | 2.8e-05 | 7.0e-03 |
| Viral mRNA Translation // Reactome |  | 1.92 | 5.1e-05 | 1.0e-02 |
| cytokine-mediated signaling pathway // BP |  | -1.92 | 2.8e-05 | 7.0e-03 |
| response to hypoxia // BP |  | -1.92 | 2.6e-05 | 7.0e-03 |
| neuropeptide signaling pathway // BP |  | 1.92 | 4.8e-04 | 4.2e-02 |
| extracellular matrix // CC |  | -1.89 | 5.1e-05 | 1.0e-02 |

### RIP140 increases GBP1 expression in BC cells

When transfected with siRIP140, MCF7 cells as well as T47D and MDA-MB-231 cells displayed a significant decrease (between 20 and 40 %) in GBP1 mRNA expression (Figure 2A), confirming the data obtained with MELN cells (Supplementary Figure 1B, left panel). Transfection of an additional siRIP140, allowed us to validate these observation (Supplementary Figure 1B, right panel). Using pro-GBP-1, a reporter construct with the GBP1 promoter driving the luciferase gene ^16^, we demonstrated that silencing RIP140 activity in MCF7, T47D and MDA-MB-231 cells reduced GBP1 promoter activity (Figure 2B). Conversely, when transfecting increasing doses of RIP140 expression plasmid, we observed that activity augmented both in MCF7 (Figure 2C) and T47D (Figure 2D) cells, indicating a transcriptional regulation of GBP1 expression by RIP140. In parallel, we also investigated the regulation of Sp100. As exhibited in Supplementary Figure 1C, a transfection of siRIP140 or siRIP140#1 also decreased Sp100 mRNA expression in MELN, MCF7 and T47D cells, while ectopic expression of RIP140 significantly augmented it (Supplementary Figure 1D). From these two examples, also consolidated by other IFNγ-regulated genes (such as *HLA-G*, *HLA-DPA1* or *IFNB1*) (data not shown), we concluded that, in various BC cells, RIP140 increased expression of IFNγ signaling pathway actors.

**Figure 2.**
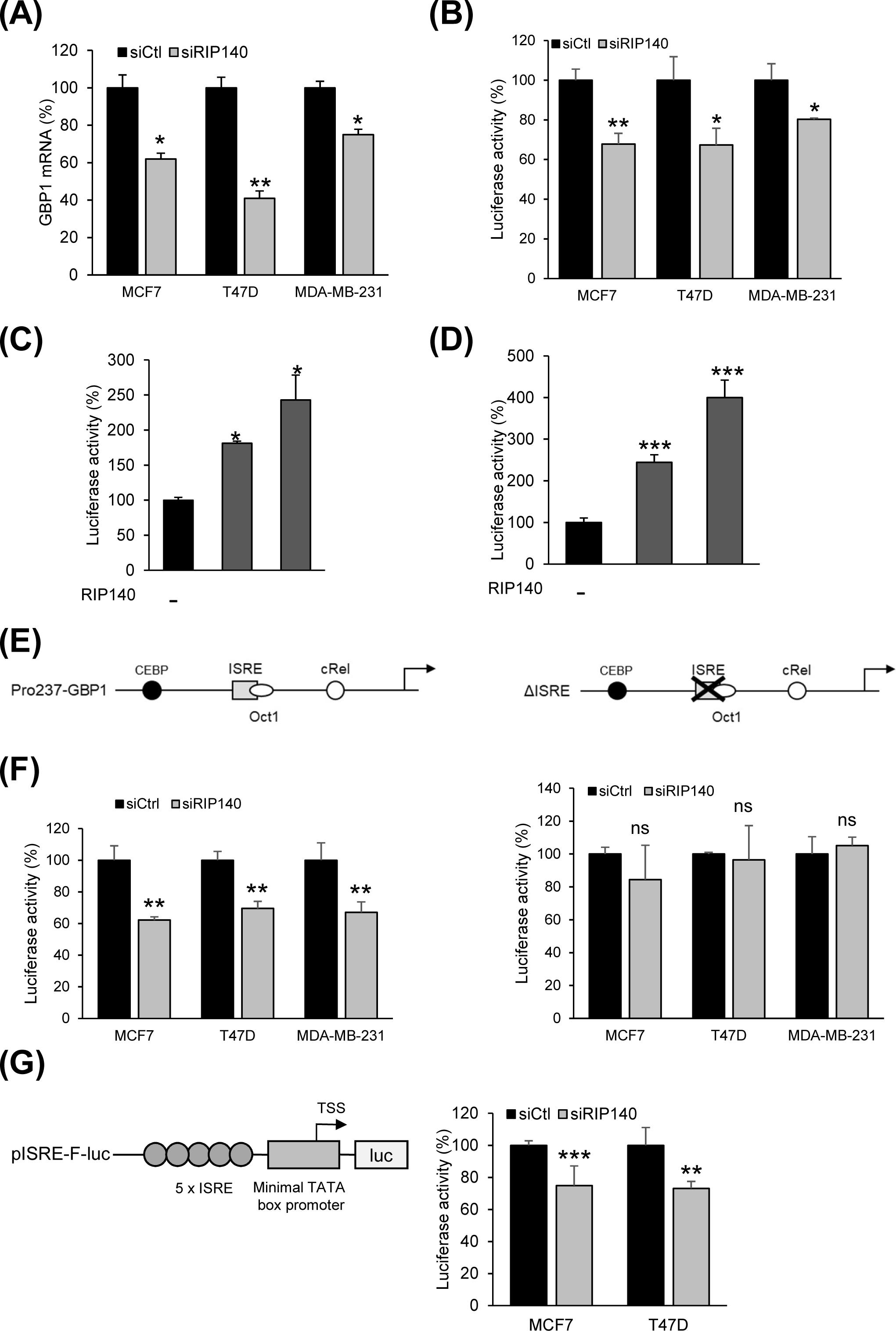
RIP140 stimulates GBP1 expression. (A) MCF7, T47D and MDA-MB-231 cells were transfected with either siCtl or siRIP140. RNAs were subject to RT-qPCR assays, and quantification of GBP1 mRNA was expressed relatively to siCtl condition. (B) MCF7, T47D and MDA-MB-231 cells were transfected with pRL-CMV-renilla and pro3757-GBP1 plasmids together with siCtl or siRIP140. Luciferase values were normalized to the renilla luciferase control and expressed relatively to the siCtl condition. MCF7 (C) and T47D (D) were transfected with pRL-CMV-renilla and pro3757-GBP1 plasmids together with increasing amounts of RIP140 expressing plasmid. Luciferase values are expressed as above described. (E) Schematic representation of pro237-GBP1 (left panel) and ΔISRE (right panel). (F) MCF7, T47D and MDA-MB-231 cells were transfected with pRL-CMV-renilla and either pro237-GBP1 (left panel) or ΔISRE (right panel) plasmids, siCtl and siRIP140. Luciferase values were normalized to the renilla luciferase control and expressed relatively to the siCtl condition. (G) Left panel: schematic representation of pISRE-F-luc. Right panel: MCF7 and T47D cells were transfected with siCtl and siRIP140 together with pISRE-F-luc and pRL-CMV-renilla. Luciferase values were expressed as in (F). Statistical analyses were performed using the Mann-Whitney test. *p<0.05, **p<0.01; ***p<0.001.

### RIP140 regulates GBP1 expression through an ISRE binding site

In order to unveil RIP140 mechanism of action on the regulation of GBP1 expression, we used a shorter version of the GBP1 promoter, *i.e.* pro237-GBP1 (Figure 2E, left panel), together with a ΔISRE mutated form, where the interferon specific response element (ISRE) is mutated ^16^ (Figure 2E, right panel). As shown in Figure 2F (left panel), transfection of siRIP140 induced a very similar decrease of pro237-GBP1 activity in MCF7, T47D and MDA-MB-231 cells as the one observed with pro-GBP-1 (Figure 2B), indicating the two constructs behaved similarly. The same result was also obtained when using siRIP140#1 (Supplementary Figure 1E, left panel). Interestingly, transfection of siRIP140 (Figure 2F, right panel) or siRIP140#1 (Supplementary Figure 1E, right panel) induced no change in ΔISRE activity revealing the importance of the response element for RIP140-dependent regulation of GBP1 promoter. Furthermore, we used a reporter plasmid consisting of five repeated ISRE sequences upstream the luciferase gene (Figure 2G, left panel). As shown in Figure 2G (right panel), both MCF7 and T47D transfected with the vector and siRIP140 displayed a significant reduction of luciferase activity. These results indicate a need for a valid ISRE to mediate RIP140 transcriptional response on the GBP1 promoter.

### RIP140 knock-down increases the IFNγ response on GBP1 expression

Upon treatment of MCF7, T47D and MDA-MB-231 cells with IFNγ (Figure 3A), expression of GBP1 mRNA was robustly stimulated (especially in T47D cells), further confirming GBP1 as a major target of IFNγ. A milder induction but still significant was also observed when the same cell lines were transfected with the pro-GBP-1 plasmid and treated with the same doses of IFNγ (data not shown).

**Figure 3.**
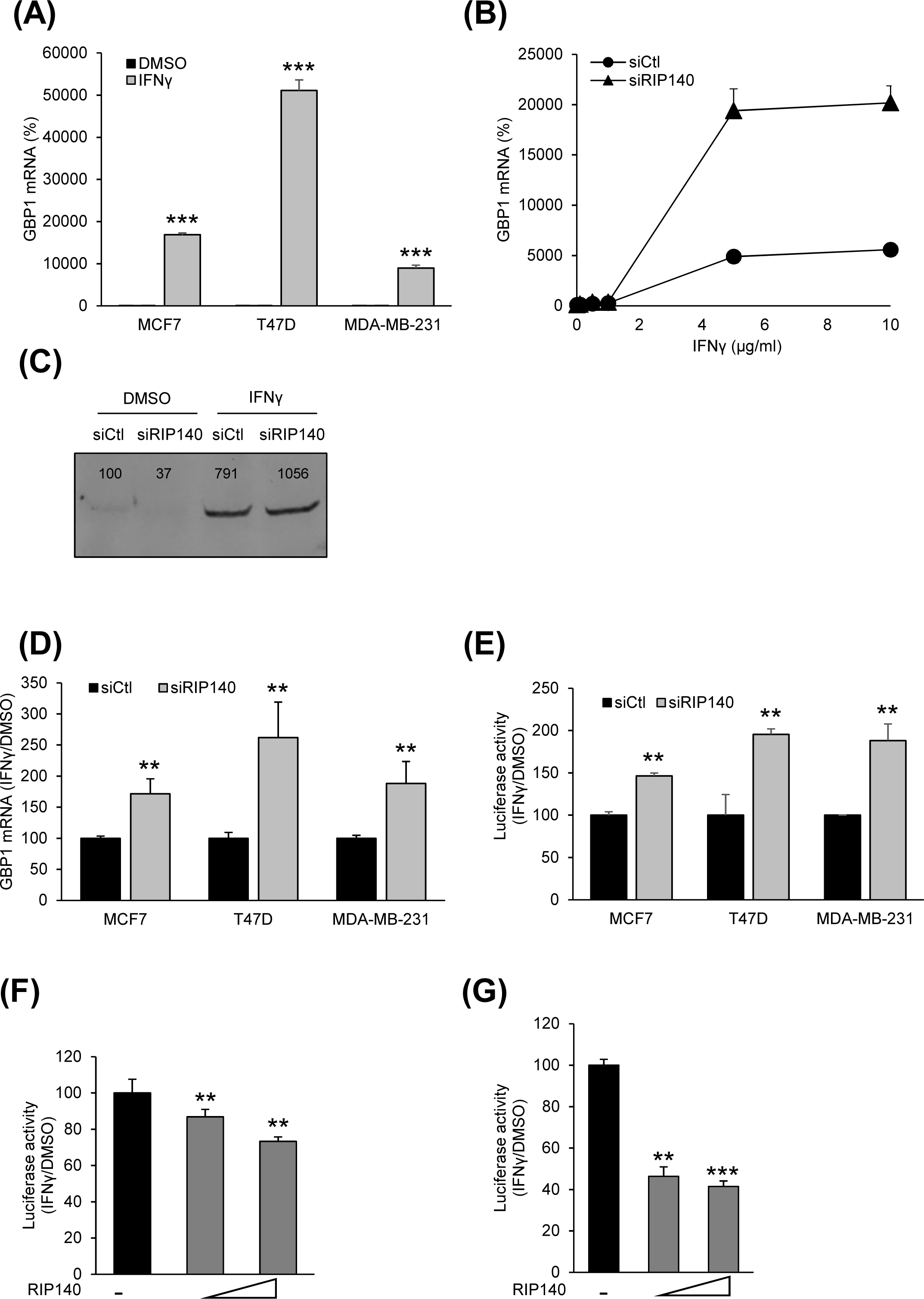
RIP140 decreases IFN-dependent GBP1 gene expression. (A) MCF7, T47D and MDA-MB-231 cells were either treated with vehicle or IFNγ (10 µg/ml) during 24h. RNAs were subjected to RT-qPCR assays, and quantification of GBP1 mRNA was expressed relatively to siCtl condition. (B) MCF7 cells were transfected with either siCtl or siRIP140 and then treated with increasing doses of IFNγ. RNAs were subject to RT-qPCR assays, and quantification of GBP1 mRNA was expressed relatively to siCtl condition. (C) MCF7 cells were transfected with either siCtl or siRIP140 and treated with either vehicle or IFNγ (10 µg/ml) during 24h. Protein extracts were submitted to western-blotting using an anti-GBP1 antibody. Relative intensity of the bands is determined by the ratio between GBP1 specific and total protein bands. (D) MCF7, T47D and MDA-MB-231 cells were transfected with either siCtl or siRIP140. Cells were then treated as in (A). GBP1 was quantified by RT-qPCR and results are expressed as the ratio between IFNγ and vehicle treatment, relatively to siCtl condition. (E) MCF7, T47D and MDA-MB-231 cells were transfected with pRL-CMV-renilla and pro3757-GBP1 plasmids. MCF7 (F) and T47D (G) cells were transfected with increasing doses of pEF-RIP140, pRL-CMV-renilla and pro3757-GBP1 plasmids. Cells (E, F and G) were treated and results expressed as in (D). Statistical analyses were performed using the Mann-Whitney test. **p<0.01; ***p<0.001.

We next wondered whether RIP140 could affect IFNγ-induced GBP1 expression. After depletion of RIP140 mRNA and treatment with increasing doses of IFNγ, we observed that the difference in GBP1 expression between siRIP140 and siCtl augmented with the dose of IFNγ in both MCF7 (Figure 3B) and T47D cells (Supplementary Figure 2A). At the protein level, while expression of GBP1 in MCF7 cells was low in basal conditions (transfection with siCtl and DMSO treatment) (Figure 3C), a decrease was observed under siRIP140 transfection. Remarkably, IFNγ induced a strong GBP1 expression, amplified when lowering RIP140 expression, confirming results obtained with mRNA. Likewise, IFNγ induction of GBP1 expression was stronger when depleting RIP140 mRNA in all three cell lines, with a stronger effect in T47D (Figure 3D and Supplementary Figure 2B). As a control, raw data obtained for T47D, in the absence and presence of IFNγ are shown in Supplementary Figure 2C. We also observed a significantly stronger IFNγ-induced activity of pro-GBP-1 under transfection of siRIP140 in MCF7, T47D and MDA-MB-231 (Figure 3E). Conversely, when ectopically expressing increasing doses of RIP140, IFNγ-induced activity of pro-GBP-1 decreased both in MCF7 (Figure 3F) and T47D (Figure 3G) cells.

In MCF7, T47D or MDA-MB231 cells transfected with RIP140, ΔISRE plasmid could no longer be stimulated by IFNγ (Supplementary Figure 2D), further demonstrating the role of the ISRE element in mediating both RIP140 and IFNγ signaling.

In parallel, focusing on Sp100 expression we showed that IFNγ significantly stimulated its mRNA expression in MCF7, T47D and MDA-MB-231 cells (Supplementary Figure 2E). We also demonstrated that IFNγ induction of Sp100 expression was stronger when transfecting siRIP140 (MCF7 and T47D cells, Supplementary Figure 2F, left panel) or siRIP140#1 (MCF7 and MDA-MB-231, Supplementary Figure 2F, right panel). This series of experiments demonstrated that low levels of RIP140 allow a full stimulatory effect of IFNγ on gene expression.

### Low levels of RIP140 increase IFNγ-driven BC cell proliferation

Based on the effect of RIP140 on IFNγ-target genes, we then wondered whether RIP140 could modulate IFNγ-dependent BC cell proliferation. Using MTT assays, we first confirmed that IFNγ inhibited the growth of MCF7 and MDA-MB-231 cells (Figure 4A, left and right panels), with a more marked effect on the latter ones. Using the same means, the two BC cell lines transfected with siRIP140 exhibited a gain in cell proliferation (Figure 4B, left and right panels), validating that RIP140 displays anti-proliferative properties in BC cells. When down-regulating RIP140 expression, we observed that IFNγ acted as a stronger repressor of either MCF7 (Figure 4C, left panel) or MDA-MB-231 (Figure 4C, right panel) cell proliferation, indicating that RIP140 prevents the cytokine to fully exert its antiproliferative effect.

**Figure 4.**
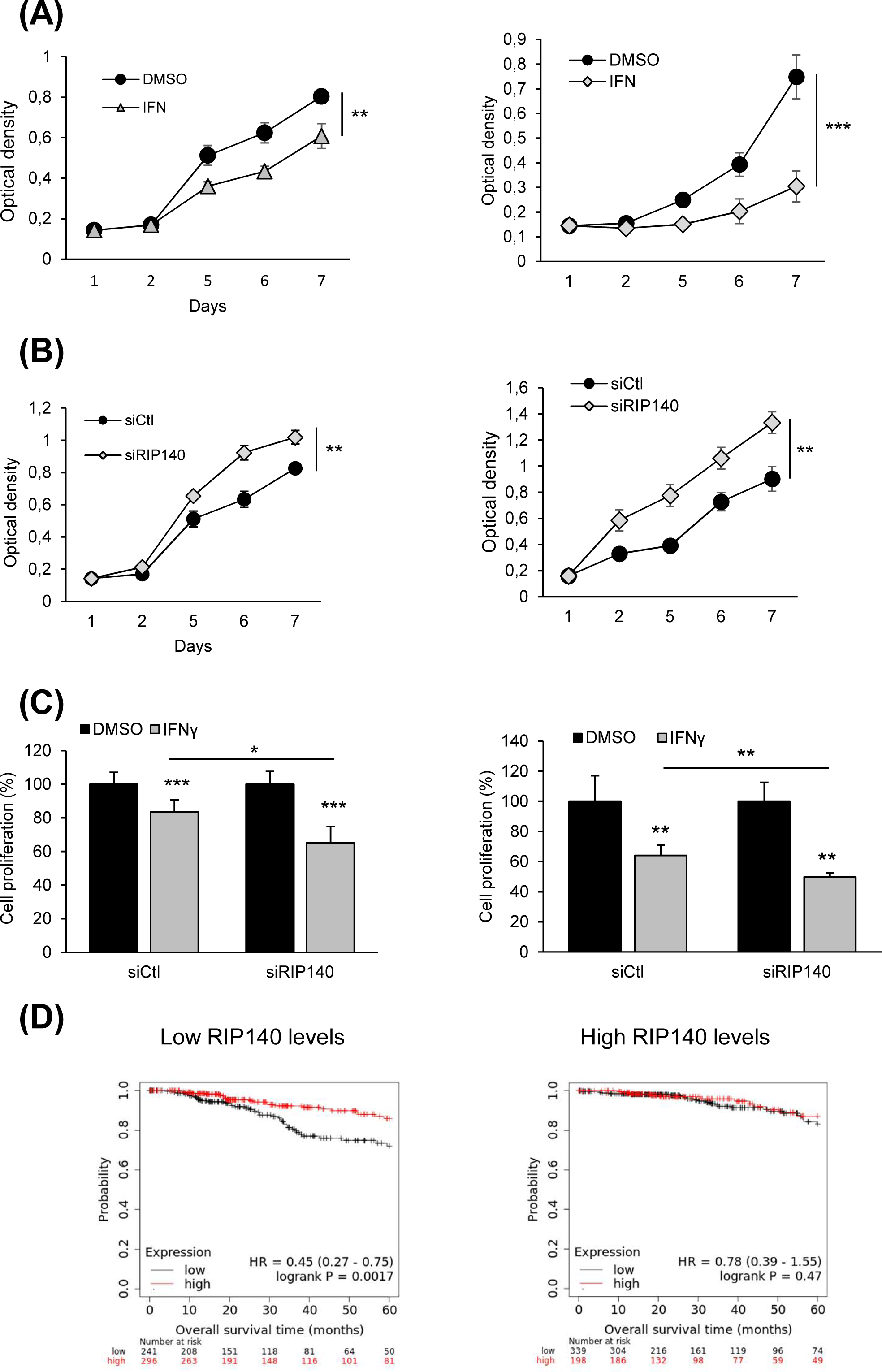
RIP140 and GBP1 regulate MCF7 and MDA-MB-231 cell proliferation. (A) MCF7 (left panel) and MDA-MB-231 (right panel) cells were treated with IFNγ (10 µg/ml) or vehicle. Cell proliferation was measured over 7 days. Absorbance of Formazan crystals was read on a spectrophotometer. (B) MCF7 (left panel) and MDA-MB-231 (right panel) cells were transfected with either siCtl or siRIP140. Cell proliferation was quantified as in (A). Cell proliferation was quantified as in (A). (C) MCF7 (left panel) and MDA-MB-231 (right panel) cells were transfected with siCtl or siRIP140. Cell proliferation was monitored as in (A) and results from day 6 were expressed relative to treatment with vehicle (DMSO). (D) The Kaplan-Meier method was used to estimate overall survival of patients from the TCGA dataset. The analysis was done on patients presenting low levels (left panel) or high levels (right panel) of *NRIP1* gene expression. Statistical analyses were performed using the Mann-Whitney test. **p<0.01; ***p<0.001.

### RIP140 expression determine the prognosis value of *IFNG* gene in BC

The experimental data shown in Fig 4C strongly suggested that high levels of RIP140 expression should inhibit the antiproliferative activity of IFNγ. We therefore hypothesized that, in breast cancers, low RIP140 levels could be a marker of such an IFNγ anti-tumor activity and consequently, that *IFNG* gene expression might be associated with an increased overall survival for patients with tumor expressing a reduced level of RIP140. Using Cox proportional hazard regression [32], we analyzed RNAseq data obtained from 1068 breast tumor samples from the TCGA dataset as described previously [33] (Figure 4). We first checked the prognostic values of the expression of each gene by analyzing patient overall survival at 60 months. High *IFNG* expression was correlated with good prognosis (HR 0.59, p=0.0011) (Supplementary Figure 3). Then we used the median as a cutoff value for classification of patients into two groups of tumors with low and high RIP140 expression, respectively. Using Kaplan-Meier plots, we investigated whether *IFNG* expression correlated with OS at 60 months within both groups using the same cutoff values. Interestingly, IFNG expression was significantly associated with increased overall survival in low RIP140 expression group (p=0.0017) but not in high RIP140 expression group (p=0.47) (Figure 4D).

## DISCUSSION

In this study, we performed a transcriptomic analysis after silencing RIP140 in MCF7 cells. We confirmed by means of qPCR, RIP140-dependent regulation of some genes, including VDR. Analysis of the sequencing data allowed us to identify IFNγ targets as being substantially modulated by RIP140. Accordingly, we identified GBP1 as a robustly RIP140-regulated gene and we described how RIP140 drives IFNγ-driven inhibition of breast cancer cell proliferation.

GBPs are a family of proteins described as IFN-inducible GTPases which can induce antimicrobial immunity and cell death ^25^. GBP1 was also associated to a bad prognosis for triple negative breast cancers ^24^. In this latter study, it must be underlined that in proliferation assays, GBP1 was shown oncogenic in MDA-MB-231 cell, whereas we described it with an anti-proliferative activity. Moreover, GBP1 expression is very heterogeneous depending on the type of cancer ^26^. It is oncogenic in prostate cancer ^27^, its expression correlates with worse survival in lung cancer ^28,29^, it is associated to treatment resistance in ovarian cancer ^30^ and overexpression of GBP1 is correlated to a poor prognosis and promotes tumor growth in glioblastoma ^31^. Conversely, GBP1 was described as a tumor suppressor in colon cancer ^32^. These observations remind of RIP140 which has a protective role in colon ^33^ or triple negative breast cancer ^12,34^ whereas a high expression is associated with stomach cancer ^35^ as well as a shorter overall survival of cervical cancer patients ^36^. GBP1 and RIP140 are a good illustration of tissue specificity of gene activity.

In this study, we described cross-talks between RIP140 and IFNγ on common transcriptional targets, mainly GBP1 and Sp100. However, we can also highlight other examples, especially E2F1 transcription factor whose expression is down-regulated by IFNγ in T47D cells ^37^ while RIP140 represses E2F1 activity ^38^. A similar cross-talk involves the androgen receptor (AR) since IFNγ is described to increase the receptor expression in prostate ^39^ while we describe a repressive activity of RIP140 on agonist-liganded AR in prostate cancer cells ^40^. We also evidenced that RIP140 was able to repress estradiol-induced AP1 activity ^41^ which was also described to induce IFNγ expression ^42^.

IFNγ is described to have anti-tumorigenic effects, noticeably by promoting apoptosis through induction of caspases expression (caspase 8 and caspase 9) ^43^. Interestingly, our RNA sequencing data together with qPCR assays evidenced a positive effect of RIP140 on the expression of the two caspases (data not shown). Alongside, IFNγ is able to repress metastasis progression though induction of fibronectin 1 (FN1) expression ^44^. Again, we also found that RIP140 was able to induce FN1 expression. Together with our cell proliferation assay, the effect of RIP140 on apoptosis and metastasis markers expression argues for a protective role of RIP140 on tumor development. By sharp contrast, low doses of IFNγ may have pro-tumorigenic effects by inducing tumor stemness through induction of ICAM1 expression ^45^. Like IFNγ, RIP140 can also induce ICAM1 (data not shown), therefore participating to a potential pro-tumorigenic effect. Interestingly, RIP140 has the same action as IFNγ on many genes, including the yin and yang effect on carcinogenesis. IFNγ also participates to cancer evasion by promoting carcinogenesis and angiogenesis. It is well characterized for papillary thyroid cancer cells where IFNγ induces epithelial-mesenchymal transition (EMT) and increases migration and invasion of the cancer cells ^46^. Even though RIP140 was not shown to promote EMT, we demonstrated that its cytoplasmic expression in breast cancer cells correlates with the expression of N-cadherin, a marker of EMT ^47^. Accordingly, together with IFNγ, RIP140 could be envisaged to participate to cancer evasion.

Finally, it has long been accepted that IFNγ plays a central role in host antitumor immunity^48^. Indeed, tumor-infiltrating lymphocytes, essential for cancer immune surveillance are the main source of IFNγ in tumors ^48^. Non-small cell lung cancer and melanoma patients treated with a PD-1 inhibitor display an increase in IFNγ protein expression accompanied with a longer progression-free survival ^49^, which could indicate the cytokine as a biomarker for prediction of response to immune checkpoint blockade. Since we describe RIP140 as a repressor of IFNγ-induced target genes, its own expression could also be hypothesized as a biomarker of the checkpoint blockade.

In conclusion, identifying IFNγ as a new target of RIP140, reinforces the major role of this transcription factor in carcinogenesis, not only within the tumor but also potentially in the tumor microenvironment.

## ACKNOWLEDGEMENTS

We would like to thank Dr Elisabeth Naschberger for the kind gift of pro3757-GBP1, pro237-GBP1 and ΔISRE plasmids as well as Dr Konstantin Maria Johannes Sparrer for the generous gift of pISRE-F-luc.

We are grateful to Dr Catherine Teyssier for the critical reading of the manuscript.

## AUTHORS CONTRIBUTIONS

SJ participated to conceptualization, formal analysis, investigation, methodology writing-original draft, AK, CP, AD, RJ and CA investigated, SS and UJ contributed to formal analysis and JC and VC participated to conceptualization, writing-original draft and funding acquisition.

## DATA AVAILABILITY STATEMENT

The data supporting the findings of our study are available from the corresponding author upon reasonable request.

## Novelty and impact

Interferon γ (IFNγ) is a major player in breast cancer. An RNA sequencing allowed to decipher the transcription factor RIP140 as a key regulator of IFNγ signaling. RIP140 represses IFNγ-dependent expression of GBP1 and modulates IFNγ-dependent repression of breast cancer cell proliferation. Finally, it was shown that IFNγ expression is associated with good prognosis for patients with low RIP140.

## Abbreviations

BC: breast cancer
GBP1: guanylate binding protein1
NRIP1: nuclear receptor interacting protein 1
RIP140: receptor-interacting protein of 140 kDa
SP100: SP100 nuclear antigen
VDR: vitamin D receptor.

