## Supplemental information for "The transcription factor RIP140 regulates interferon γ signaling in breast cancer"

| siRNA | Sequence |
|---|---|
| siCtl | UAAUGUAUUGGAACGCAUA |
| siRIP140 | GAAGCGUGCUAACGAUAAA |
| siRIP140#1 | AUACGAAUCUUCCUGAUGU |
| siGBP1 | AAGGCAUGUACCAUAAGCU |

| Primer name | Forward | Reverse |
|---|---|---|
| RIP140 | AATGTGCACTTGAGCCATGATG | TCGGACACTGGTAAGGCAGG |
| ACBD5 | CCAAAGAGGAAGCCATGATT | GACACGCAGCAATTCTTCAA |
| MAEA | CAGTTGTCCATGACCCTGAA | TCTCCCGGTCAATGTTCTTC |
| VDR | TGGCTTTCACTTCAATGCTATGA | CGTCGGTTGTCCTTGGTGAT |
| GBP1 | AGGAGTTCCTTCAAAGATGTGGA | GCAACTGGACCCTGTCGTT |
| SP100 | TCCATGACAAATTGCCTCTCC | GAGATGGGGAACCCGAAGG |

**Supplementary table 1.** List of the different siRNAs and primers used in this study for qPCR quantification together with their forward and reverse sequences.

**Supplementary table 1**

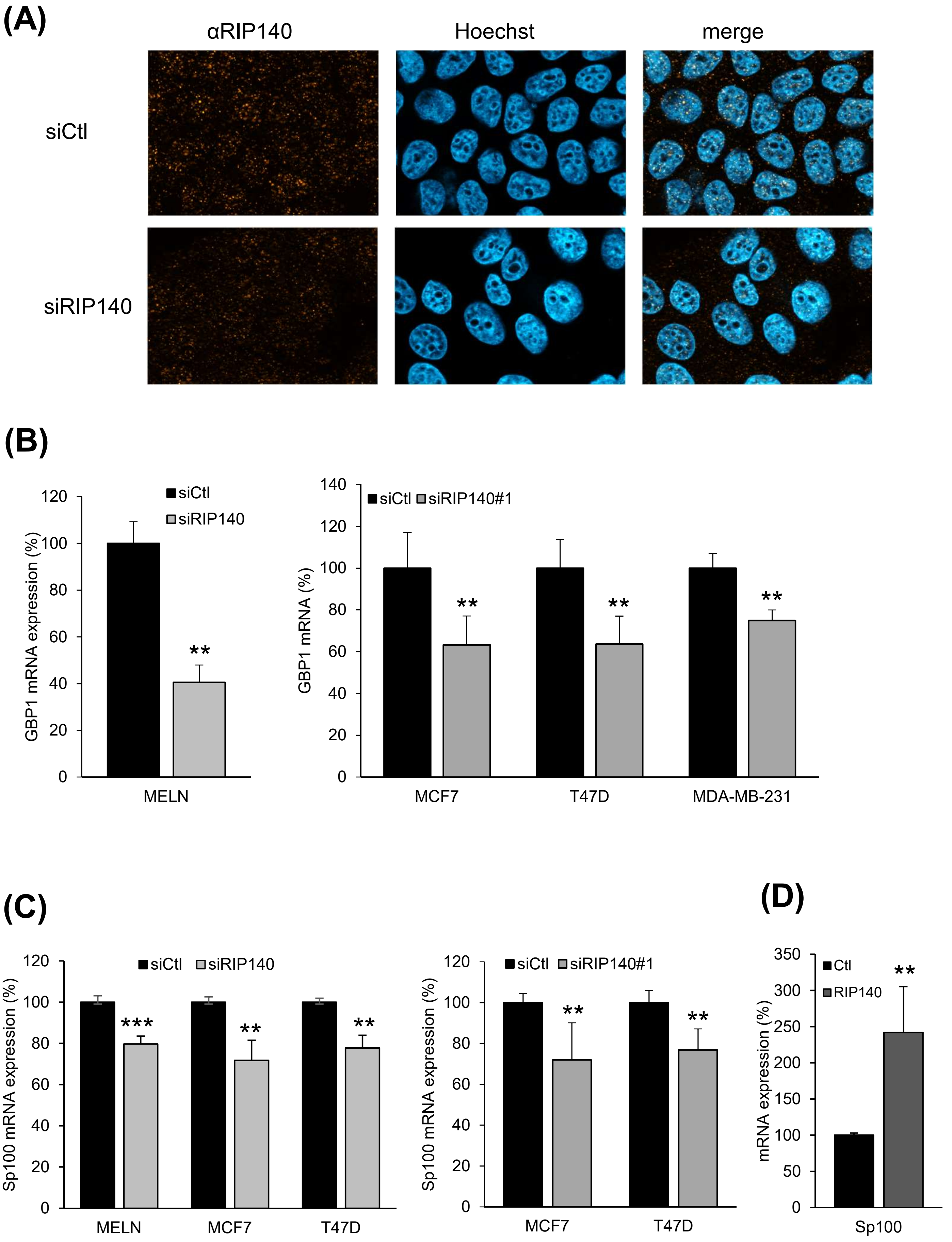


**Supplementary Figure 1**

**(E)**

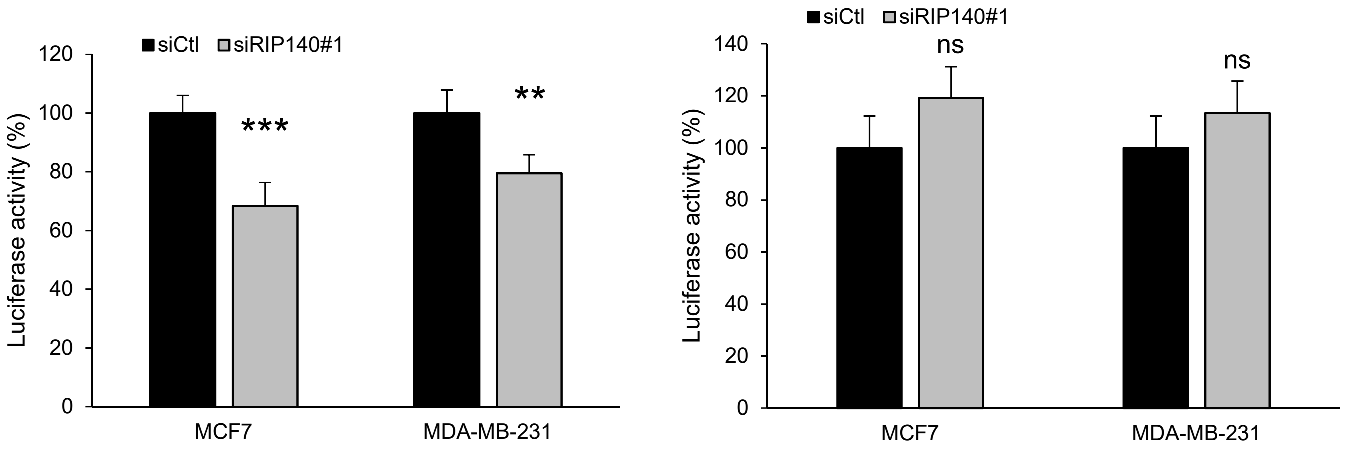


**Supplementary Figure 1.** (A) MCF7 cells were analyzed by immunofluorescence after transfection of siCtl (control) or siRIP140, using anti-RIP140 antibody (ab42126, Abcam) and Hoechst labeling. Images were acquired with a Zeiss Imager M2 microscope with the Apotome system and a Plan Apochromat 40×/1.3 DIC (oil) controlled by the ZEN software (Carl Zeiss). (B) MELN (left panel) or MCF7, T47D and MDA-MB-231 (right panel) cells were transfected with either siCtl, siRIP140 or siRIP140#1 as indicated. RNAs were subjected to RT-qPCR assays, and quantification of GBP1 mRNA was expressed relatively to siCtl condition. (C) MELN, MCF7 and T47D were transfected with either siCtl, siRIP140 or siRIP140#1 as indicated and Sp100 mRNA was quantified and expressed as in A. (D) MCF7 cells were transfected with either pEF-RIP140 (RIP140) or the empty vector (Ctl). Sp100 mRNA was quantified and expressed as in (B). (E) MCF7 and MDA-MB-231 cells were transfected with pRL-CMV-renilla and either pro237-GBP1 (left panel) or ΔISRE (right panel) plasmids, siCtl and siRIP140#1. Luciferase values were normalized to the renilla luciferase control and expressed relatively to the siCtl condition. Statistical analyses were performed using the Mann-Whitney test. **p<0.01; ***p<0.001.

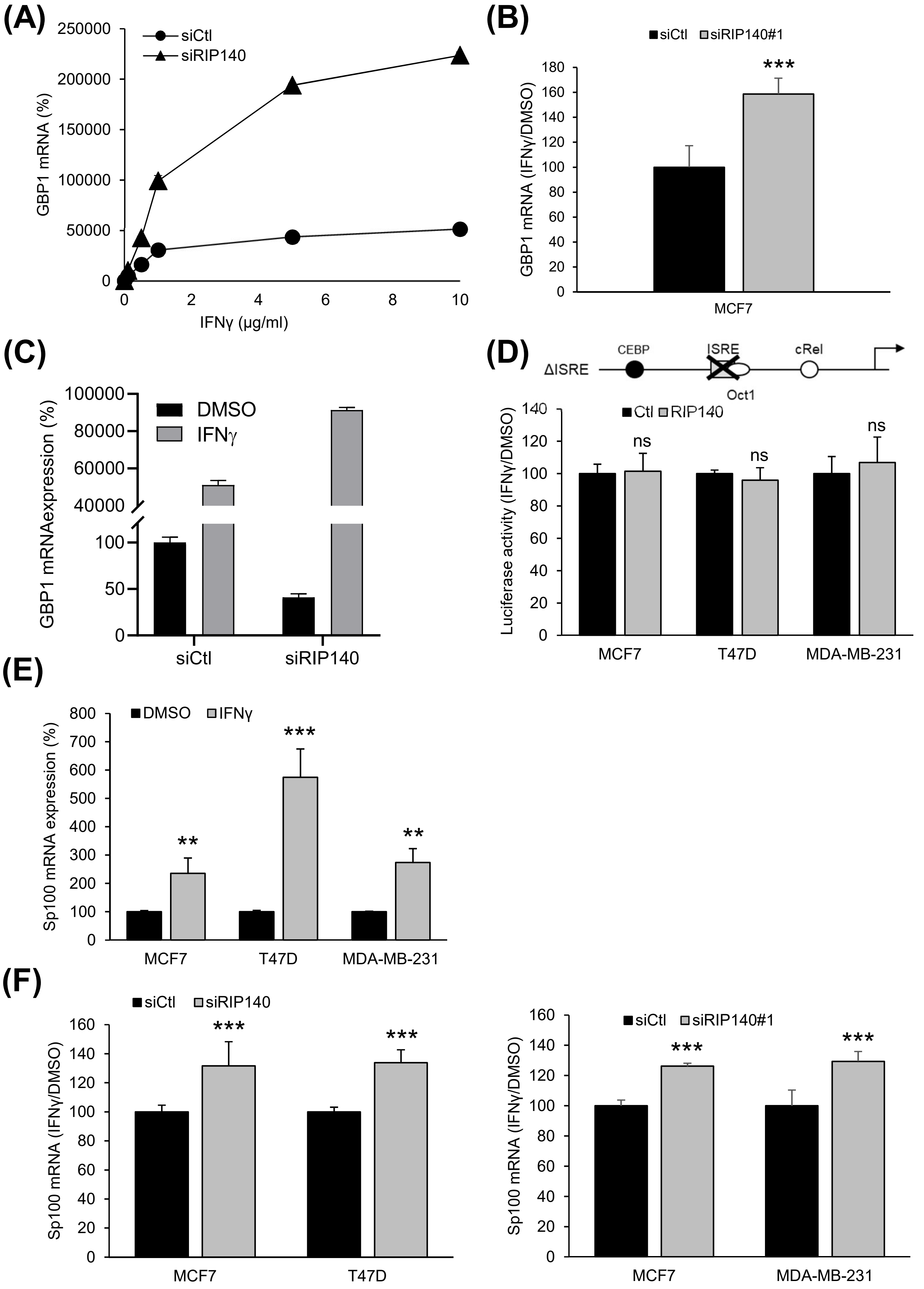


**Supplementary Figure 2**

**Supplementary Figure 2.** (A) T47D cells were transfected with either siCtl or siRIP140 and then treated with increasing doses of IFNγ. RNAs were subject to RT-qPCR assays, and quantification of GBP1 mRNA was expressed relatively to siCtl condition.
(B) MCF7 cells were transfected with siCtl or siRIP140#1 and treated with IFNγ (10 µg/ml) or vehicle during 24h. GBP1 was quantified by RT-qPCR and results are expressed as the ratio between IFNγ and vehicle treatment, relatively to siCtl condition.
(C) T47D cells were transfected with either siCtl or siRIP140 and then treated or not with IFNγ (10 µg/ml). Quantification of GBP1 mRNA was done as in (B). (D) Schematic representation of ΔISRE (upper panel). MCF7, T47D and MDA-MB-231 (lower panels) cells were transfected with pRL-CMV-renilla, Ctl or RIP140 and ΔISRE plasmids and then treated with either vehicle or IFNγ (10 µg/ml) during 24h. Luciferase activity was expressed as the ratio between IFNγ and vehicle treatment, relatively to siCtl condition.
(E) MCF7, T47D and MDA-MB-231 cells were either treated with IFNγ (10 µg/ml) or vehicle during 24h. Sp100 quantification was performed as in A. (F) MCF7 T47D and MDA-MB-231 cells were transfected with siCtl, siRIP140 or siRIP140#1 and treated with IFNγ (10 µg/ml) or vehicle during 24h treated. Sp100 was quantified as in (A). Statistical analyses were performed using the Mann-Whitney test. **$p<0.01$; ***$p<0.001$.

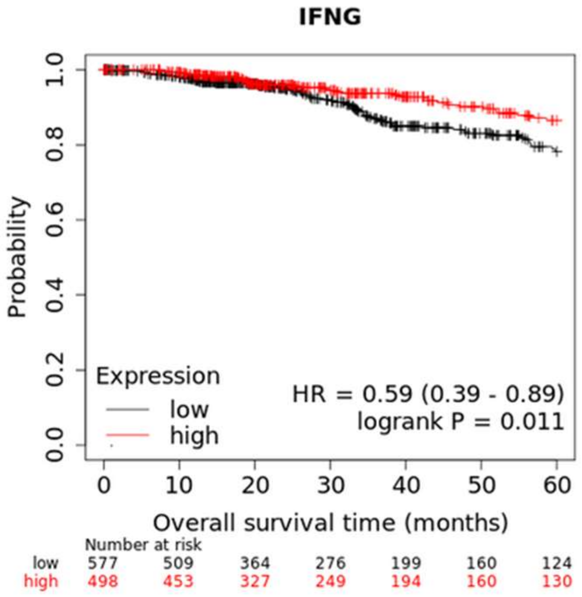


**Supplementary Figure 3.** The Kaplan-Meier method was used to estimate overall survival of patients from the TCGA dataset. The analysis was performed according to *IFNG* gene expression levels.

**Supplementary Figure 3**
